# EM-net: Deep learning for electron microscopy image segmentation

**DOI:** 10.1101/2020.02.03.933127

**Authors:** Afshin Khadangi, Thomas Boudier, Vijay Rajagopal

## Abstract

Recent high-throughput electron microscopy techniques such as focused ion-beam scanning electron microscopy (FIB-SEM) provide thousands of serial sections which assist the biologists in studying sub-cellular structures at high resolution and large volume. Low contrast of such images hinder image segmentation and 3D visualisation of these datasets. With recent advances in computer vision and deep learning, such datasets can be segmented and reconstructed in 3D with greater ease and speed than with previous approaches. However, these methods still rely on thousands of ground-truth samples for training and electron microscopy datasets require significant amounts of time for carefully curated manual annotations. We address these bottlenecks with EM-net, a scalable deep convolutional neural network for EM image segmentation. We have evaluated EM-net using two datasets, one of which belongs to an ongoing competition on EM stack segmentation since 2012. We show that EM-net variants achieve better performances than current deep learning methods using small- and medium-sized ground-truth datasets. We also show that the ensemble of top EM-net base classifiers outperforms other methods across a wide variety of evaluation metrics.

## I. Introduction

Among a variety of biological microscopy imaging modalities, segmenting three-dimensional (3D) EM stacks is challenging [1]. Such datasets are notoriously difficult to work with since the texture and intensity variations across all parts of the images are similar [2, 3]. A variety of mechanisms have been proposed to address this problem using machine-learning approaches, including ilastik [4], TrakEM2 [5] and Microscopy Image Browser [6]. Although such tools demonstrate acceptable inference metrics, deep learning offers a further advancement which involves the adaptation of the underlying parameters (features) to the data automatically [7, 8].

Several deep learning network models have been successfully used for segmenting electron microscopy images. For example, U-net [9] extended the architecture of Fully Convolutional Networks (FCNs) [10] and utilised up-sampling in opposite expansive path with skip connections to obtain the output mask within an end-to-end learning framework. It has been widely used by both medical and biological image analysis communities and is still regarded as one of thenoise most successful models in this context. CDeep3M, a Plug-and-Play cloud-based Deep Convolutional Neural network (DCNN) [11] was proposed to segment electron microscopy images, as well as X-ray images. The proposed model involves the ensemble of 3 different classifiers which processes the data with multiple numbers of image frames, i.e. 1, 3 and 5 image mini-batches. The model has been implemented using a variety of datasets including microCT X-ray and fluorescence microscopy, SBF-SEM and EM stacks.

Most of the deep learning methods used for EM image analysis utilise the network topologies proposed previously on ImageNet challenge [12] as an encoder [13-15]. A decoder is subsequently used to retrieve the original resolution. Such methods depend on large ground-truth samples, however, according to one study [16], manual data segmentation or annotation costs $10 per μm^3^ on average for EM volumes, which might incur thousands of dollars for a relatively large data volume. Additionally, providing ground-truth samples for EM datasets is challenging due to their inherently low contrast. Moreover, the evaluation metrics for EM image segmentation reported in the literature remain sparse, and currently available methods have not been benchmarked. The computational efficiency of existing deep learning methods for EM analysis are also yet to be investigated.

In this study, we propose EM-net, a scalable deep neural network for rapid learning from limited ground-truth samples for 2D EM image segmentation. In this new network architecture, we propose trainable rectifiers as activation functions to capture intricate nonlinear structures within EM image volumes with minimal computational complexity. EM-net demonstrates reliable performance when trained based on both small- and medium-sized ground-truth datasets. Moreover, we benchmark segmentation metrics for EM image segmentation and show that EM-net base classifiers provide better results when compared to existing methods despite their low computational complexities. Finally, we show that an ensemble of top base classifiers based on EM-net significantly outperform the previously proposed methods.

## II. METHOD

### A. Trainable Linear Unit (TLU)

We propose an adaptive activation function “TLU” which introduces two additional trainable parameters per activation function along with other parameters of the network. We implemented the proposed activation function on TensorFlow [17], and a user can initialise these parameters with desired distributions, impose constraints, utilise regularisation and even share these parameters across the feature channels. Figure 1 illustrates the TLU where *α* and *β* represent the adaptive parameters, as shown in Equation 1 (here we have implicitly treated tan *α* and tan *β* as *α* and *β*, respectively):

**Fig. 1.**
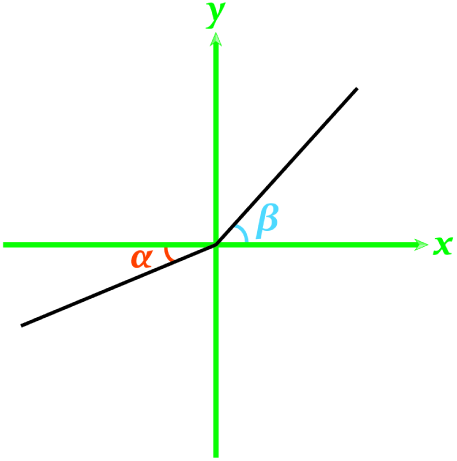
Illustration of the Trainable Linear Unit activation function. α and β represent the adaptive parameters for negative and positive values, respectively.

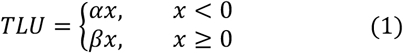

#### 1) Definition

Consider the TLU defined in the format of activation function as follows:

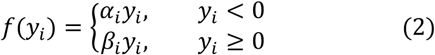

where *y*_*i*_ is the input to the nonlinear activation function *f* on the *i*th channel, *α*_*i*_ and *β*_*i*_ are the coefficients controlling the slopes of the activation function. The subscript *i* implies that TLU can be varied on different channels. When *α*_*i*_ = 0 and *β*_*i*_ = 1, TLU becomes ReLU; when *α*_*i*_ ≠ 0 and *β*_*i*_ = 1, it becomes Parametric ReLU (PReLU) [18]. Equation 2 is equivalent to *f*(*y*_*i*_) = *β*_*i*_ max(0, *y*_*i*_) + *α*_*i*_ min(0, *y*_*i*_). If *α*_*i*_ is small and constant and *β*_*i*_ = 1, TLU becomes Leaky ReLU (LReLU) [19]. The main motivation of Leaky ReLU was to avoid zero gradients. It had been shown that Leaky ReLU provides an insignificant improvement on accuracy compared to ReLU. Moreover, PReLU demonstrates sensitivity to initialisation as investigated comprehensively in [18]. On the contrary, TLU generalises ReLU, LReLU and PReLU, and provides better inference results as shown in subsequent sections.

TLU introduces only a few extra parameters relative to the size of the other network parameters, so the risk of overfitting is minimal. Moreover, the channel-wide shared variant only adds two additional parameters per feature channel. Hence, the added time of training will also be negligible.

#### 2) Optimisation

Similar to PReLU, TLU can be trained using backpropagation [20] and optimised concurrently with other parameters. We employ the chain rule to derive the update formula. The gradients of *α*_*i*_ and *β*_*i*_ for one layer are as follows:

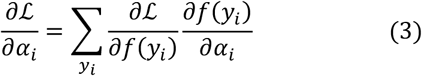

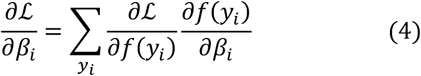

where ℒ represents the loss function. The first term 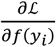 is the gradient propagated from the deeper layer. The gradient of the activation function is defined as 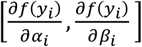 where:

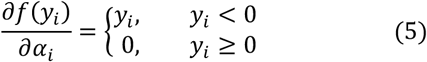

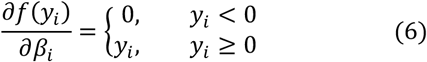

The summation 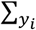 is performed over the entire components of the feature map. For the channel-shared variant, the gradients are obtained by 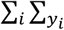, where ∑_*i*_ runs over all channels of the layer.

To update the *α*_*i*_ and *β*_*i*_, the momentum method can be obtained as follows:

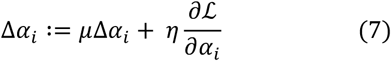

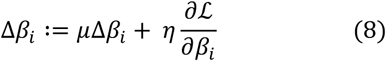

Where *μ* and *η* represent momentum and learning rate, respectively. We do not use regularisation or constraints over the *α*_*i*_ and *β*_*i*_, and we use *α*_*i*_ = 0 and *β*_*i*_ = 1 for initialisation.

### B. Network topology

U-net [9] has been widely used in many applications, especially in medical image analysis. We adopt the same concept of the encoder-decoder scheme, as illustrated in figures 2 – 3. The proposed network “EM-net” is comprised of cells which possess unique properties and fixed arrangement. The primary motivation behind the architecture design was to furnish the network with skip connections in block levels rather than only in encoder-decoder level as it has been shown that skip connections are vital to biomedical image segmentation [21]. All these processing units of the EM-net are equipped with TLUs and come within two different structures, as shown in the figures. The first type of these cells comprises three layers of two-dimensional convolutions each having the same number of feature channels where first and third layers are merged to form the output of the cell (coloured in orange, figures 2 – 3). However, the second type includes four layers of two-dimensional convolutions, where the odd layers have twice the number of feature channels of the even layers. Moreover, the second and fourth layers are merged to form the output in these cells (coloured in purple, figures 2 – 3).

**Fig. 2.**
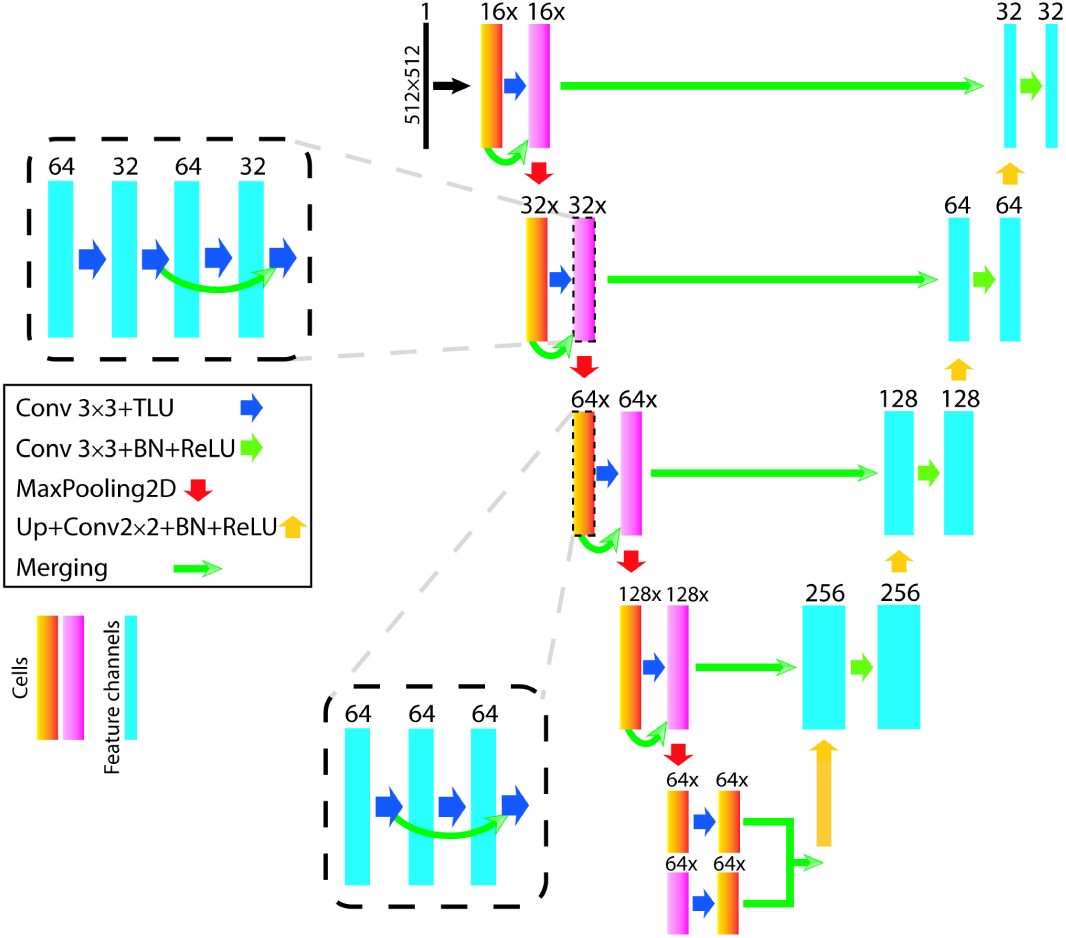
network topology of EM-net V1 2X. Orange and purple represent the cells and blue represents 2D convolutional feature channels. TLUs are used in the encoder and Rectified Linear Units (ReLUs) are employed in the decoder. Note the merging between layers in cells and between cells in the encoder. Similar to U-net, the outputs of each block in the encoder are merged with the same resolution block in the decoder.

**Fig. 3.**
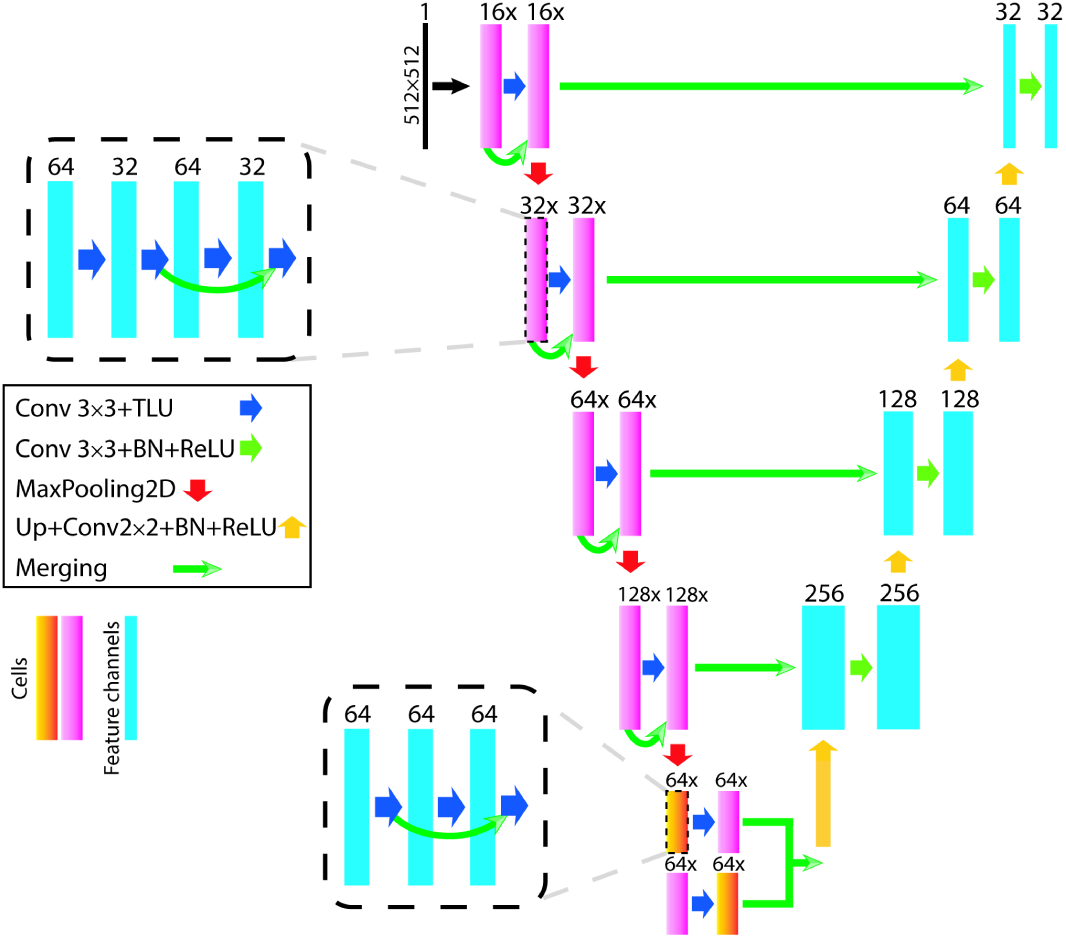
network topology of EM-net V2. Orange and purple represent the cells and blue represents 2D convolutional feature channels. TLUs are used in the encoder and Rectified Linear Units (ReLUs) are employed in the decoder. Note the merging between layers in cells and between the cells, and the differences in the structures of bottlenecks between figures 2 and 3.

EM-net mainly involves two types of variants as part of the ablation studies. Figure 2 represents the architecture of “EM-net V1 2X” in which “V1” stands for the first variant, and “2X” means that the width of this network is twice the width of “EM-net V1 BN” where “BN” implies that batch-normalization has been utilised in the encoder in addition to the decoder. All the EM-net variants use batch-normalization [22] in the decoder, and they exhibit simpler structure in the decoders or expansive paths. The first variant of EM-net is structured by the orange and purple cell types, as shown in figure 2. However, the second variant of the EM-net is dominated by second type cell except for the network bottleneck in which two different types of cells interact with each other as illustrated in figure 3. We conducted ablation studies on a variety of scaled versions of EM-net in terms of width and depth. Hence, “EM-net V1 BN”, “EM-net V1 BN 2X”, “EM-net V1 2X”, “EM-net V1 4X”, “EM-net V2”, “EM-net V2 2X” and “EM-net V2 4X” were proposed, each having 5.4m, 6.8m, 7m, 28m, 7.5m, 14.8m, and 33.8m parameters, respectively. Details of all these networks have been provided in tutorials on GitHub^1^. Here only “EM-net V1 2X” and “EM-net V2” are shown in figures 2 and 3, respectively.

## III. EXPERIMENTS

We performed experiments to validate the proposed method. We participated in the ISBI 2012 challenge on EM stacks segmentation for neuronal structures [23] and also applied our method to a FIB-SEM dataset to segment mitochondria in cardiomyocytes [24]. We compared the performance of EM-net with CDeep3M [11], U-net [9], VGG-16 [25], ResNet-50 [26], SegNet [27] and PReLU-net [18]. We utilised a wide spectrum of metrics for evaluation purposes including F1-score, Foreground-restricted Rand Scoring after border thinning (*V*^*Rand*^_(*thinned*)_) Foreground-restricted Information-Theoretic Scoring after border thinning (*V*^*Info*^_(*thinned*)_) [23], accuracy, sensitivity, specificity, positive and negative predictive values and Jaccard similarity index. We also compared the convergence times of these methods to address one of our main aims which is rapid learning from limited ground-truth data. Finally, we report the behaviours of these classifiers during the training in terms of training and validation accuracy, F1-score, loss and AUC-ROC [28].

### A. Data

We evaluated the proposed method using two publicly available datasets. The first dataset was obtained from the ISBI 2012 EM segmentation challenge [23]. The training dataset is a stack of 30 slices from Serial Section Electron Microscopy images of Drosophila first instar larva ventral nerve cord [29], measured at 2×2×1.5 microns with the resolution of 4×4×50 nm/voxel. We split the data randomly into training and validation sets by 70% and 30%, respectively. We augmented the training data using shifting, zooming, rotation, flipping, mirroring and shearing. No elastic distortion was implemented during the augmentation. The augmented dataset had the same resolution as original dataset (512×512 pixels). The test dataset on this competition has the same size as the training dataset which is 30×512×512.

The second dataset includes left ventricular myocyte FIB-SEM datasets collected from three mice, as described previously [30]. We applied the same augmentation techniques as described for the neuronal dataset and extracted 24 random patches each having 512×512 pixels for training, validation and testing purposes. After we manually annotated mitochondria on this sample, we split the data randomly into training, validation and testing by ^16^/_24_, ^4^/_24_ and ^4^/_24_, respectively.

### B. Training

We used Rectified Adam (RAdam) optimiser [31] as the main optimisation method for both datasets. The learning rate for both datasets was set to 0.001, and we did not use weight decay during our implementations. We specifically chose RAdam for optimisation because of the several advantages it offers. First, Rectified Adam exhibits much more robust behaviour to learning rate variations as it rectifies the variance of the learning rate. Secondly, it provides better generalisation on a variety of datasets compared to other state-of-the-art optimisation techniques [31]. We used binary cross-entropy as the loss function for both datasets. Using spatial dropout layers [32] improved our results. The training and validation batch size was set to 4 for both experiments. All the experiments except CDeep3M were implemented using TensorFlow GPU 1.8.0 CUDA 9.0 [17] and KERAS 2.2.4 [33]. All the experiments were performed on a GPU cluster, HPC Spartan [34] with 4 NVIDIA Tesla P100 on each node. We parallelised all the implementations on one node, utilising 4 GPU devices during training, validation and testing.

For the implementation of CDeep3M, we used Amazon Web Services (AWS) and launched a stack of 100GB GPU instance on p3.2xlarge in the US West (Oregon) region. CDeep3M is implemented in Caffe, and we used the default settings for training as described in [11]. We set the maximum iterations to 30,000. Since our data was 2D, we used one frame classifier on CDeep3M.

### C. The initialisation of convolutional layers for TLU

We initialised the convolutional layers with He Normal [18] and Glorot Normal [35] and obtained similar results which imply that TLU is less sensitive to the initialisation of convolutional layers as compared with PReLU. We used He Normal throughout our implementations in this study.

## IV. RESULTS

### A. ISBI 2012 challenge for EM stack segmentation on neuronal structures

We submitted the outputs of the EM-net to ISBI 2012 challenge on EM stack segmentation after the training. We did not perform any post-processing or segmentation refinement, and we submitted the outputs of the network to the challenge directly. “EM-net V1 2X” achieved 1^st^ place on this challenge among all the submissions that did not utilise segmentation refinement. The network achieved 8^th^ place in overall scoring. We fine-tuned the last ten layers of the network by training the network after freezing the remaining layers. Table 1 represents the results of our submissions to the ISBI challenge. The full leader board can be accessed on the challenge website.

**TABLE I.** Illustration of results from leaderboard of ISBI 2012 challenge on EM stacks segmentation. EM-net achieved first place among those methods that did not utilise post-processing algorithms. The overall score of EM-net on the ISBI leader board was eighth and was achieved with only 7 million parameters.

| Method (no post-process) | $\nu_{Rand}$ | $\nu_{Info}$ |
| --- | --- | --- |
| EM-net | **0.9836** | 0.9872 |
| ACE-Net [36] | 0.9797 | 0.9898 |
| M2FCN [37] | 0.9780 | 0.9902 |
| FusionNet [14] | 0.9780 | 0.9899 |
| U-net [9] | 0.9727 | 0.9866 |

### B. EM-net variants achieve lowest validation loss and highest F1-score

We visualised the behaviour of all these networks, excluding CDeep3M during training by tracking validation accuracy, loss, F1-score, and AUC-ROC. As shown in figure 4, EM-net variants achieve the lowest validation loss and highest F1-score. Moreover, this figure implies that using batch-normalization boosts the validation AUC-ROC and F1-score as EM-net V1 BN and BN 2X have achieved the desirable scores in these metrics.

**Fig. 4.**
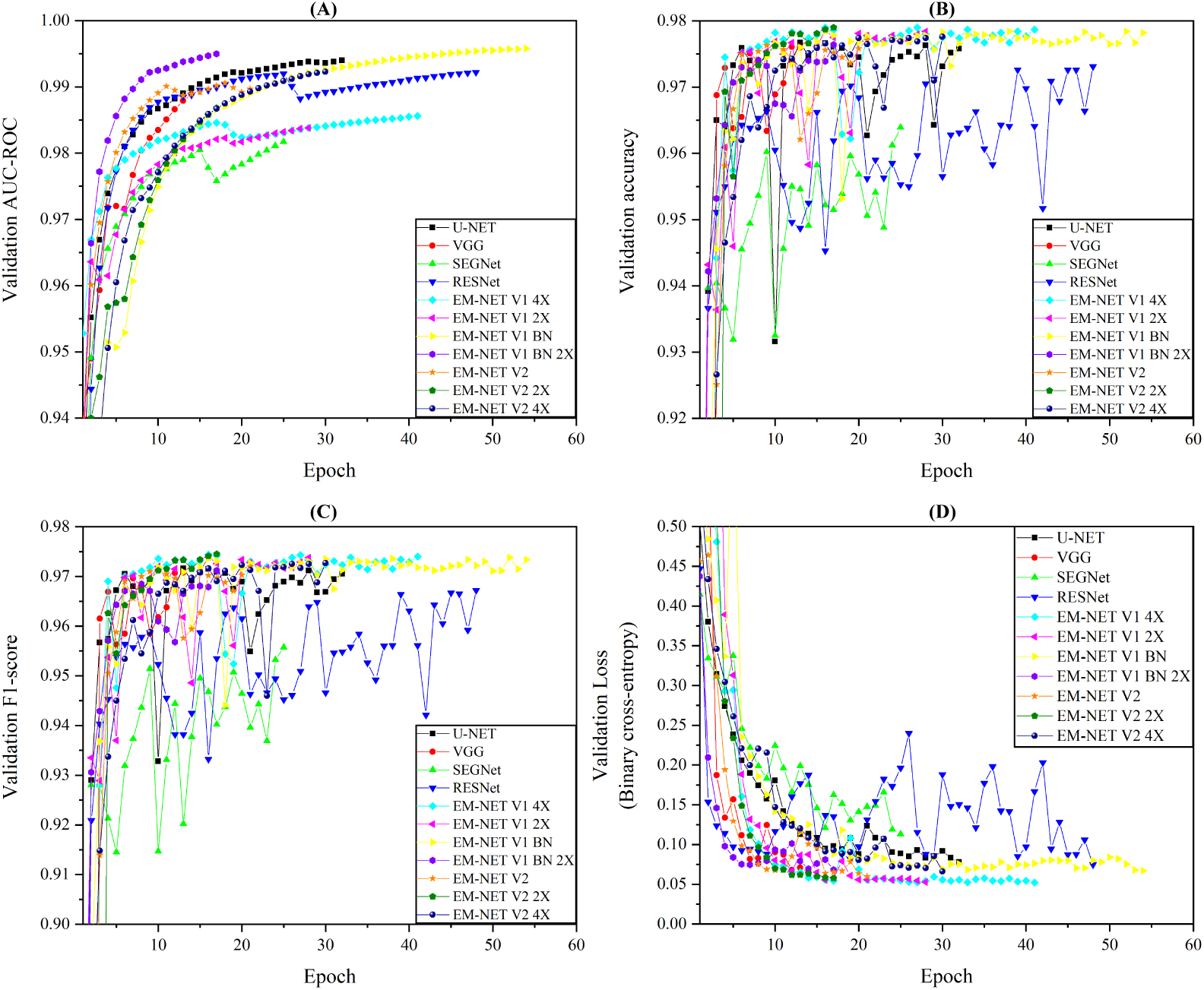
Illustration of the validation AUC-ROC (A), accuracy (B), F1-score (C) and loss (D) for different methods during training for mitochondria segmentation on the cardiac dataset. As shown, EM-net variants have achieved the best validation convergence losses and using batch-normalization in encoder has boosted validation AUC-ROC.

### C. EM-net converges faster and has less complexity in terms of FLOPS

One of the other main aims of this study was to develop a method that demands less computational resources while converging in minimal time and achieving desirable performances. We investigated the performance of the EM-net in the previous section and showed that EM-net outperforms existing methods on segmenting electron microscopy data. Hence, we dedicate this section to investigate the complexity and convergence times of different methods. We show that EM-net converges faster or demonstrates comparable convergence times. Moreover, we report the complexity of these networks in terms of floating-point operations per second (FLOPS) to compare their requirement for computational resources. We did not include the convergence time of CDeep3M here as the convergence time for this method took 15 hours in total on AWS. The convergence time for the other techniques are reported in terms of hours and based on several experiments.

Figure 5 represents the convergence times on the cardiac dataset based on our experiments using four Tesla P100 GPUs. According to our investigations, top-performing EM-net base classifiers (V1 2X, V1 BN 2X, V2, V2 2X and V2 4X) demonstrate faster or comparable convergence times compared to other methods. As shown, EM-net demonstrates less variability in convergence times which indicates that its computational performance is robust regardless of the network scale. However, other methods show greater variability, which implies that they are less robust and much more sensitive to initialisation.

**Fig. 5.**
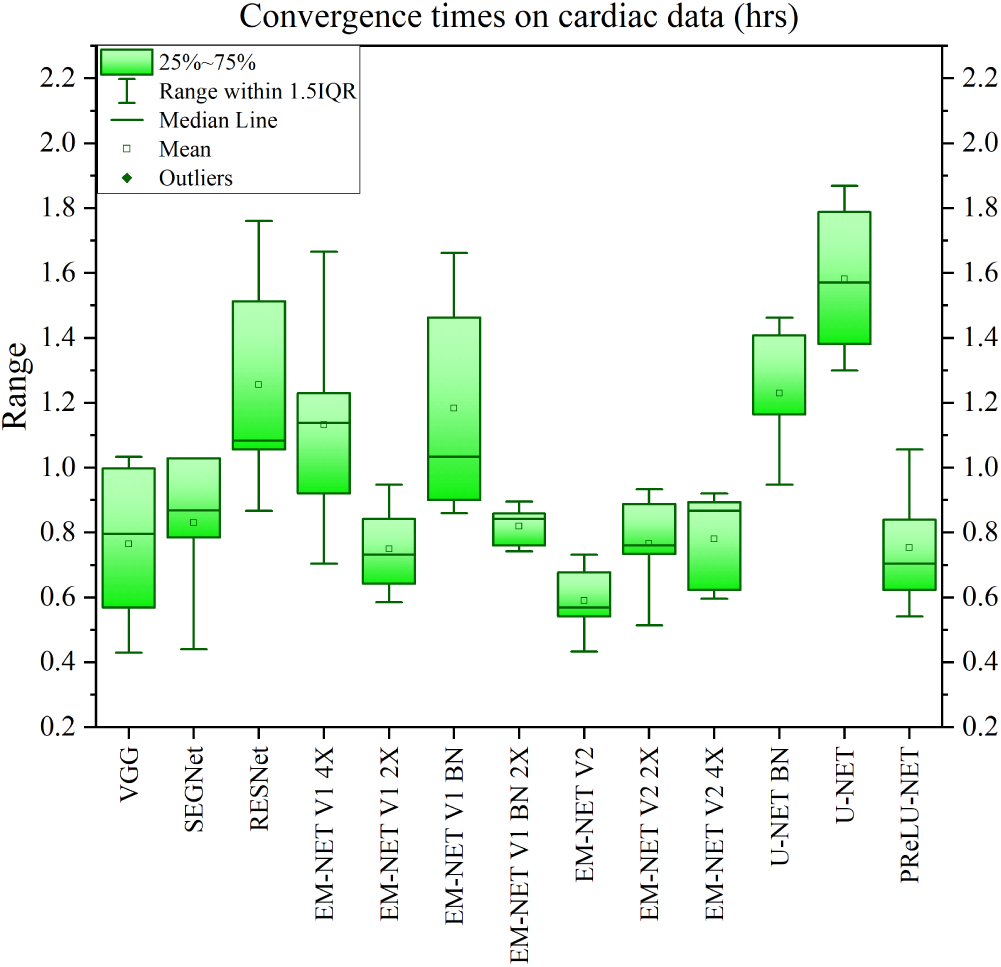
Illustration of the convergence times on cardiac dataset based on our experiments using four Tesla P100 GPUs. As shown, top EM-net base classifiers (EM-net V1 2X, V1 BN 2X, V2, V2 2X and V2 4X) demonstrate less or comparable convergence times compared to other methods.

Figure 6 illustrates the number of FLOPS, number of parameters and *V*^*Rand*^_(*thinned*)_ score of the corresponding methods. These reports were generated on TensorFlow, and the input tensor had a shape of (1, 512, 512, 1) representing a single batch of a single-channel image with a resolution of 512×512 pixels. As shown in this figure, ResNet and CDeep3M required the lowest and highest FLOPS, respectively. However, EM-net required 200 billion FLOPS on average, including all base classifiers, and 150 billion FLOPS on average when excluding “V1 4X”. That is 40% more efficient than the average of other methods which is 346 billion FLOPS. Hence, EM-net will require less computational resources compared to other methods needed for a single forward pass. As shown, EM-net variants achieve highest *V*^*Rand*^_(*thinned*)_ scores with low FLOPS and number of parameters.

**Fig. 6.**
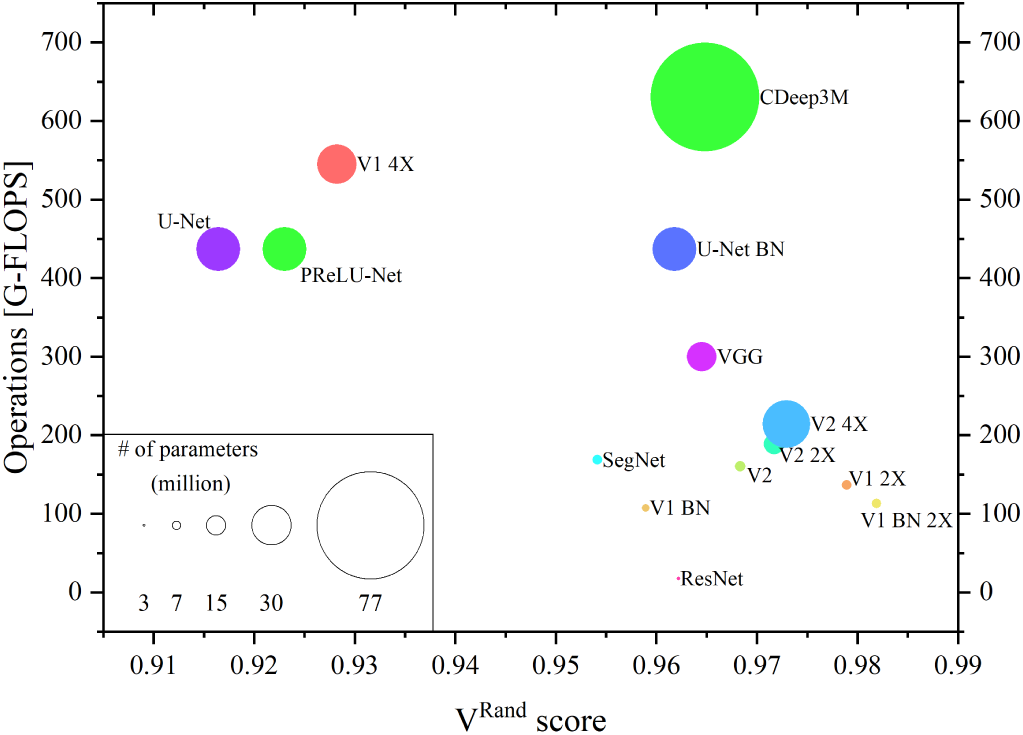
ball chart reporting the *V*^*Rand*^_(*thinned*)_ score vs. computational complexity in terms of FLOPS. EM-net base classifiers achieve highest *V*^*Rand*^_(*thinned*)_ scores while demonstrating low computational complexity and a small number of parameters. CDeep3M and ResNet show the highest and lowest number of parameters and computational complexity, respectively.

### D. The ensemble of top EM-net base classifiers outperforms the existing methods

We used average and majority voting to obtain an ensemble of top EM-net base classifiers based on F1-score, *V*^*Info*^_(*thinned*)_ and *V*^*Rand*^_(*thinned*)_. The results are shown in Table 2 and Figure 7. Table 2 represents the evaluation results on the test dataset for segmenting mitochondria on cardiac data.

**TABLE II.** Comparison of results on segmenting mitochondria using the cardiac test dataset. The ensemble of the top 5 EM-net base classifiers outperforms other methods in terms of evaluation metrics investigated in this study. BN, PPV and NPV represent batch-normalisation, positive predictive value and negative predictive value, respectively.

| Method | Module | F1-score | $V_{Rand}$ | $V_{Info}$ | Accuracy | Sensitivity | Specificity | PPV | NPV | Jaccard |
| --- | --- | --- | --- | --- | --- | --- | --- | --- | --- | --- |
| CDEEP3M | - | 0.9795 | 0.96485 | 0.96845 | 0.9745 | 0.9638 | 0.993 | 0.9958 | 0.9405 | 0.9344 |
| VGG | - | 0.9801 | 0.96452 | 0.96347 | 0.9745 | 0.9889 | 0.9494 | 0.9714 | 0.9802 | 0.9315 |
| SEGNET | - | 0.9815 | 0.95414 | 0.95408 | 0.9764 | 0.9864 | 0.9592 | 0.9767 | 0.9759 | 0.937 |
| RESNET | - | 0.9785 | 0.96219 | 0.96593 | 0.9729 | 0.9735 | 0.9717 | 0.9836 | 0.9548 | 0.929 |
| EMNET | V1 4X | 0.9846 | 0.92822 | 0.95021 | 0.9804 | 0.9894 | 0.9647 | 0.9798 | 0.9813 | 0.9472 |
| EMNET | V1 2X | 0.9852 | 0.97893 | 0.97674 | 0.9811 | 0.99 | 0.9657 | 0.9804 | 0.9823 | 0.9491 |
| EMNET | V1 BN | 0.9854 | 0.95894 | 0.96294 | 0.9815 | 0.9867 | 0.9725 | 0.9842 | 0.9769 | 0.9506 |
| EMNET | V1 BN 2X | 0.9788 | 0.98188 | 0.97739 | 0.9799 | 0.9854 | 0.9561 | 0.9723 | 0.9768 | 0.946 |
| EMNET | V2 | 0.9801 | 0.96833 | 0.96057 | 0.9745 | 0.9911 | 0.9457 | 0.9694 | 0.984 | 0.9313 |
| EMNET | V2 2X | 0.9842 | 0.97172 | 0.96854 | 0.9798 | 0.989 | 0.9639 | 0.9794 | 0.9806 | 0.9459 |
| EMNET | V2 4X | 0.9849 | 0.97294 | 0.97439 | 0.9808 | 0.9892 | 0.9661 | 0.9806 | 0.981 | 0.9484 |
| U-NET | BN | 0.9811 | 0.96182 | 0.94929 | 0.9758 | 0.9875 | 0.9554 | 0.9747 | 0.9778 | 0.9352 |
| U-NET | - | 0.9611 | 0.91642 | 0.93644 | 0.9521 | 0.9713 | 0.922 | 0.9511 | 0.9536 | 0.8824 |
| PReLU-NET | - | 0.9745 | 0.92302 | 0.94799 | 0.9688 | 0.9789 | 0.953 | 0.9702 | 0.9665 | 0.9225 |
| ENSEMBLE | Majority Voting | 0.9869 | 0.98728 | 0.98803 | 0.9833 | 0.991 | 0.97 | 0.9828 | 0.9842 | 0.9551 |
| ENSEMBLE | Average Voting | 0.9872 | 0.98692 | 0.98792 | 0.9839 | 0.985 | 0.982 | 0.9895 | 0.9743 | 0.9572 |

**Fig. 7.**
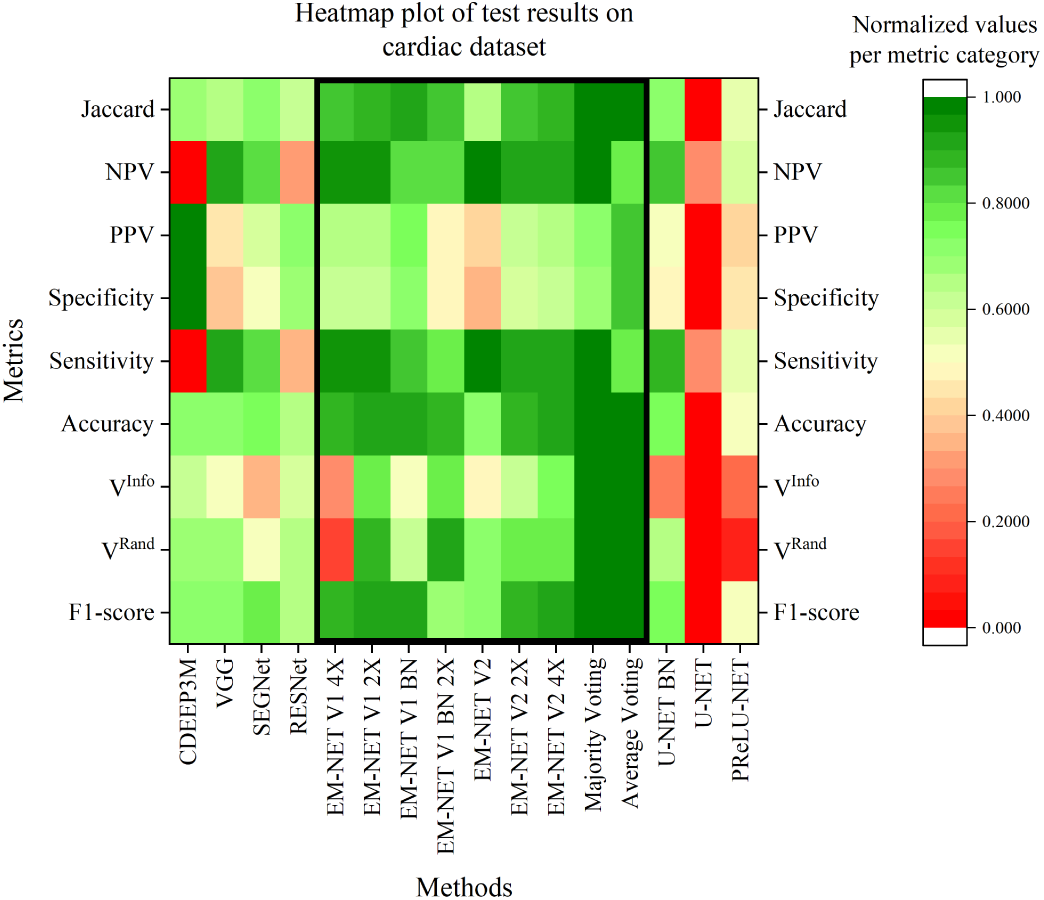
Heatmap of evaluation metrics for different methods based on the cardiac test dataset. This heatmap is, in fact, the distilled version of performance tables; however, the values are normalised using min-max normalisation per metric category to favour the ranking criteria. The black box shows the grouping of EM-net family, and as shown, the ensemble of top EMNET base classifiers using average or majority voting boost the segmentation performance with great magnitude.

As shown, the majority of EM-net base classifiers outperform other methods based on the above evaluation metrics, which we consider as main evaluation metrics in this study. In addition to the above, we chose the top 5 base classifiers and obtained an ensemble of them using average and majority voting. The results imply that the ensemble of EM-net base classifiers outperforms the existing methods based on the majority of evaluation metrics, especially the first three metrics.

Figure 7 represents the heatmap of test results along with the evaluation metrics. To rank the segmentation performances and help the reader to compare the results shown in table 2, we have used min-max normalisation to normalise the results for each of the evaluation metrics. The performances of EM-net are demonstrated within a bounded black box on this figure. As shown, EM-net base classifiers achieve better results compared to other methods as the majority of green patches are distributed within the box mentioned above. Moreover, the ensemble of EM-net classifiers (top 5) outperforms other methods as they have achieved the maximum evaluation metric values (the greenest patches are limited to average and majority voting).

### E. Visualisation of intermediate feature channels reveals the superiority of TLU over ReLU

To measure the effect of TLU on learnt features in deep neural networks, we visualised the intermediate feature channels of two networks with the same topology but different activation functions in encoders. We chose EM-net V1 2X for this purpose and used ReLU in all the layers of the encoder in the first network and utilised TLU in the second. The results are shown in Figure 8. We randomly chose three layers, including two activation layers and one max-pooling layer. As shown, most of the features in the first network where we have used ReLU instead of TLU are irrelevant and sparse. However, the feature channels extracted in the second network demonstrate unique and meaningful features and the number of sparse feature channels is minimal. That is because TLU enforces individual feature channels to adapt to data variations and exhibit specialised distributions unique to different feature channels. Moreover, as shown in Figure 8, the usage of ReLU has resulted in some of the feature channels to become sparse as the gradients for those feature channels have stopped adapting to the data.

**Fig. 8.**
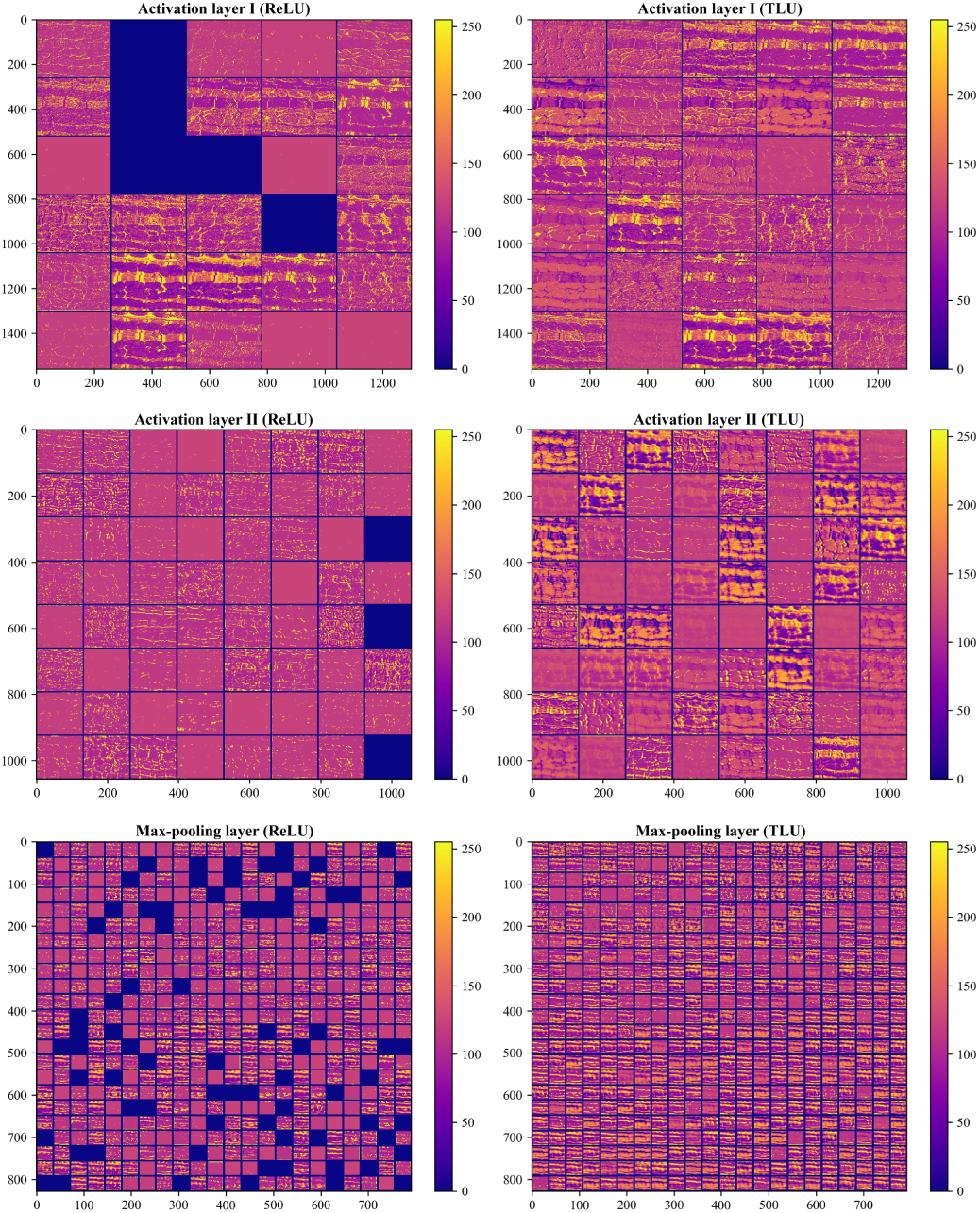
Visualisation of intermediate feature channels reveals that TLU assists the extraction of more meaningful features and minimises the risk of fitting to redundant and sparse features. Left column: ReLU, Right column: TLU.

### F. Visualisation of segmentation results on test data imply EM-net is less prone to false positives

We visualised the segmentation results on the cardiac test dataset by overlaying the binary segmentations maps on ground-truth masks. Figure 9 represents the results of this investigation where yellow, green, red and black correspond to true positive, false positive, false negative (missing mitochondria) and true negative, respectively. We found that EM-net variants are less prone to false positive errors as compared to all other methods. Moreover, as shown, missing mitochondria or false negatives are significantly minimised in EM-net based on an ensemble of average and majority voting when compared to base classifiers. CDeep3M is more prone to false positives and produces minimal false negatives or missing mitochondria. VGG and U-net BN are more prone to missing mitochondria.

**Fig. 9.**
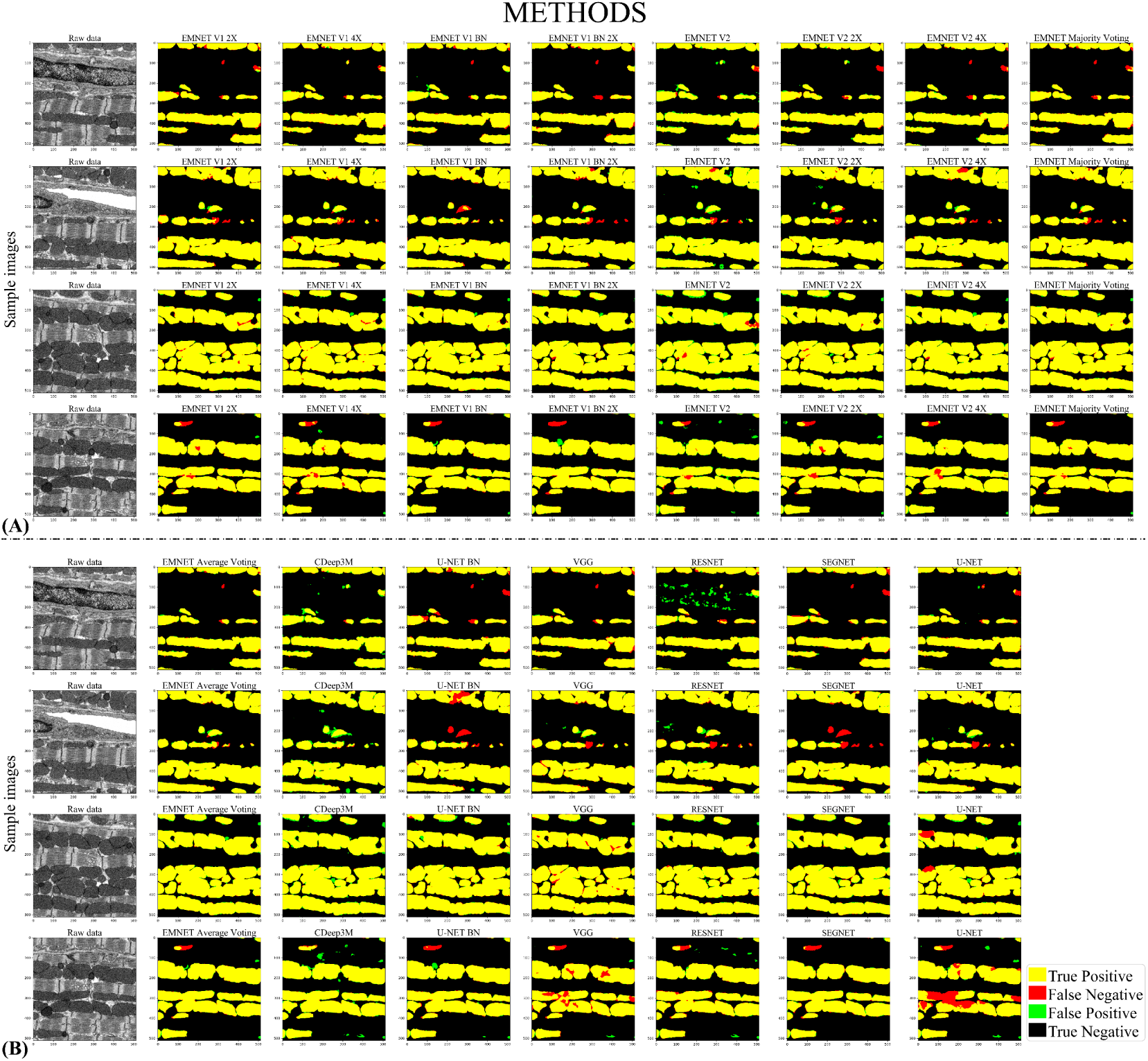
Comparison of segmentation results on cardiac test data. Yellow, green, red and black correspond to true positive, false positive, false negative (missing mitochondria) and true negative, respectively. EM-net variants are less prone to false positives. Using ensembles further minimises false negatives. Top 4 rows (A) represents the results for all of the EM-net base classifiers along with the results of the ensemble method using majority voting. Bottom 4 rows (B) illustrates the results for other methods along with the ensemble of EM-net top-performing base classifiers using average voting.

## V. CONCLUSIONS

In this paper, we proposed EM-net, a DCNN which utilises newly introduced trainable rectifiers to capture intricate nonlinearities in EM data. EM-net is scalable and demonstrates competitive performance over the other methods despite low computational complexity. We benchmarked the evaluation metrics and showed that EM-net base classifiers achieve better results, where an ensemble of these classifiers outperform the existing methods with great magnitude. We provide the full implementation on TensorFlow with the pre-trained network models. We believe that EM-net applications can be easily extended to other tasks, including 3D volume segmentation and we are quite confident that proposed TLU would provide good results for other imaging modalities.

## Acknowledgement

This research was undertaken using the LIEF HPC-GPGPU Facility hosted at the University of Melbourne. This Facility was established with the assistance of LIEF Grant LE170100200.

## Footnotes

1 https://github.com/akhadangi

